# Highly accurate and robust protein sequence design with CarbonDesign

**DOI:** 10.1101/2023.08.07.552204

**Authors:** Milong Ren, Chungong Yu, Dongbo Bu, Haicang Zhang

**Author notes:** Corresponding authors: D.B. and H.Z. (, lead contact).

## Abstract

Protein sequence design, the inverse problem of protein structure prediction, plays a crucial role in protein engineering. Although recent deep learning-based methods have shown promising advancements, achieving accurate and robust protein sequence design remains an ongoing challenge. Here, we present CarbonDesign, a new approach that draws inspiration from successful ingredients of AlphaFold for protein structure prediction and makes significant and novel developments tailored specifically for protein sequence design. At its core, CarbonDesign explores Inverseformer, a novel network architecture adapted from AlphaFold’s Evoformer, to learn representations from backbone structures and an amortized Markov Random Fields model for sequence decoding. Moreover, we incorporate other essential AlphaFold concepts into CarbonDesign: an end-to-end network recycling technique to leverage evolutionary constraints in protein language models and a multi-task learning technique to generate side chain structures corresponding to the designed sequences. Through rigorous evaluations on independent testing data sets, including the CAMEO and recent CASP15 data sets, as well as the predicted structures from AlphaFold, we show that CarbonDesign outperforms other published methods, achieving high accuracy in sequence generation. Moreover, it exhibits superior performance on *de novo* backbone structures obtained from recent diffusion generative models such as RFdiffusion and FrameDiff, highlighting its potential for enhancing *de novo* protein design. Notably, CarbonDesign also supports zero-shot prediction of the functional effects of sequence variants, indicating its potential application in directed evolution-based design. In summary, our results illustrate CarbonDesign’s accurate and robust performance in protein sequence design, making it a promising tool for applications in bioengineering.

## Main

Protein sequence design, also referred to as inverse protein folding, is to identify amino acid sequences that can fold into a given protein backbone structure while exhibiting desired functions. It serves as a crucial step in computational protein design, which has recently made significant advancements in the engineering of therapeutics [1, 2], enzymes [3, 4], and more applications [5]. Typically in *de novo* protein design, determining the optimal sequences became essential once the backbone structures are derived from either energy-based methods [6] or recent diffusion generative models [7–9].

Recent advancements in deep learning-based sequence design methods have demonstrated promising results in generating highly accurate candidate sequences [10–15]. These approaches differ from one another in their strategies for encoding the protein structure and decoding the associated sequences. Typically, ProteinMPNN [10] and ESM-IF [11] utilize neural networks to encode the entire backbone structure and subsequently decode the sequences in an end-to-end autoregressive manner. On the other hand, methods such as 3DCNN [12], ABACUS-R [13], and ProDESIGN-LE [14] individually encode the structural context of each residue and iteratively refine the designed sequences, starting from a randomly initialized sequence.

Protein structure prediction and protein sequence design are closely intertwined, with advancements in one field benefiting the other. Inspired by the remarkable success of AlaphFold2 [16] and RosettaFold [17] in addressing the protein folding problem, we adapt their key concepts to the inverse folding and propose a novel approach CarbonDeisgn, aiming to improve sequence design through enhancing the encoder and decoder architecture, leveraging more efficient features, and refining the training strategy.

At its core, CarbonDesign utilizes a novel network architecture Inverseformer to transform 3D structural features into single and pair representations using a series of node updates and triangular edge updates, following a Markov Random Field (MRF) module for sequence decoding. Intuitively, Inverseformer inverts the information flow compared to AlphaFold’s Evoformer and primarily focuses on learning representations from backbone structures.

We also introduce two other crucial concepts. Firstly, we adopt the network recycling strategy [16, 18, 19] to recycle the entire network with shared weights in an end-to-end manner. During the recycling stages, we incorporate sequence embedding from the protein language model EMS2 [20], enabling CarbonDesign to fuse evolutionary and structural constraints effectively. Second, we leverage multi-task learning with several auxiliary losses to guide the learning of single and pair representations directly and to predict the sequences and the corresponding side chain structures.

We extensively evaluate CarbonDesign using diverse datasets, including the CAMEO dataset [21], the recently released CASP15 dataset [22], and the predicted structures from AlphaFold. Additionally, in the context of *de novo* protein design, we further assess the utility of CarbonDesigin in reconstructing sequence for the *de novo* structures derived from diffusion generative methods such as RFdiffusion [7] and FrameDiff [8]. Furthermore, we demonstrate that CarbonDesgin serves as a reliable zero-shot predictor of mutational effects on protein function, with its performance evaluated using deep mutational scanning datasets encompassing millions of missense variants.

## Results

### Model architecture

CarbonDesign improves protein sequence design by incorporating novel neural network architectures and training procedures based on evolutionary and structural constraints. To convert protein 3D structure to 1D sequence, we invert and adapt the network architecture employed in AlphaFold, which was initially developed for 3D structure prediction from 1D sequence (Figure 1 and Table 1).

**Table 1:**
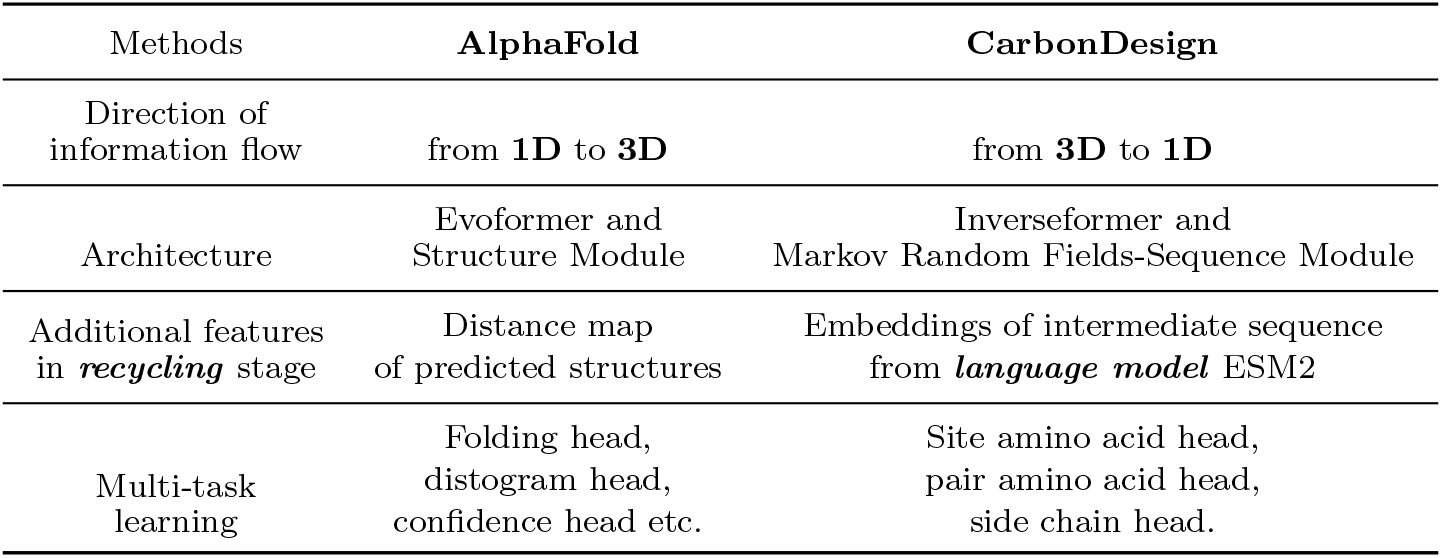
Key concepts of CarbonDesign inspired by AlphaFold.

| Methods | AlphaFold | CarbonDesign |
| --- | --- | --- |
| Direction of information flow | from **1D** to **3D** | from **3D** to **1D** |
| Architecture | Evoformer and Structure Module | Inverseformer and Markov Random Fields-Sequence Module |
| Additional features in *recycling* stage | Distance map of predicted structures | Embeddings of intermediate sequence from *language model* ESM2 |
| Multi-task learning | Folding head, distogram head, confidence head etc. | Site amino acid head, pair amino acid head, side chain head. |

**Fig. 1:**
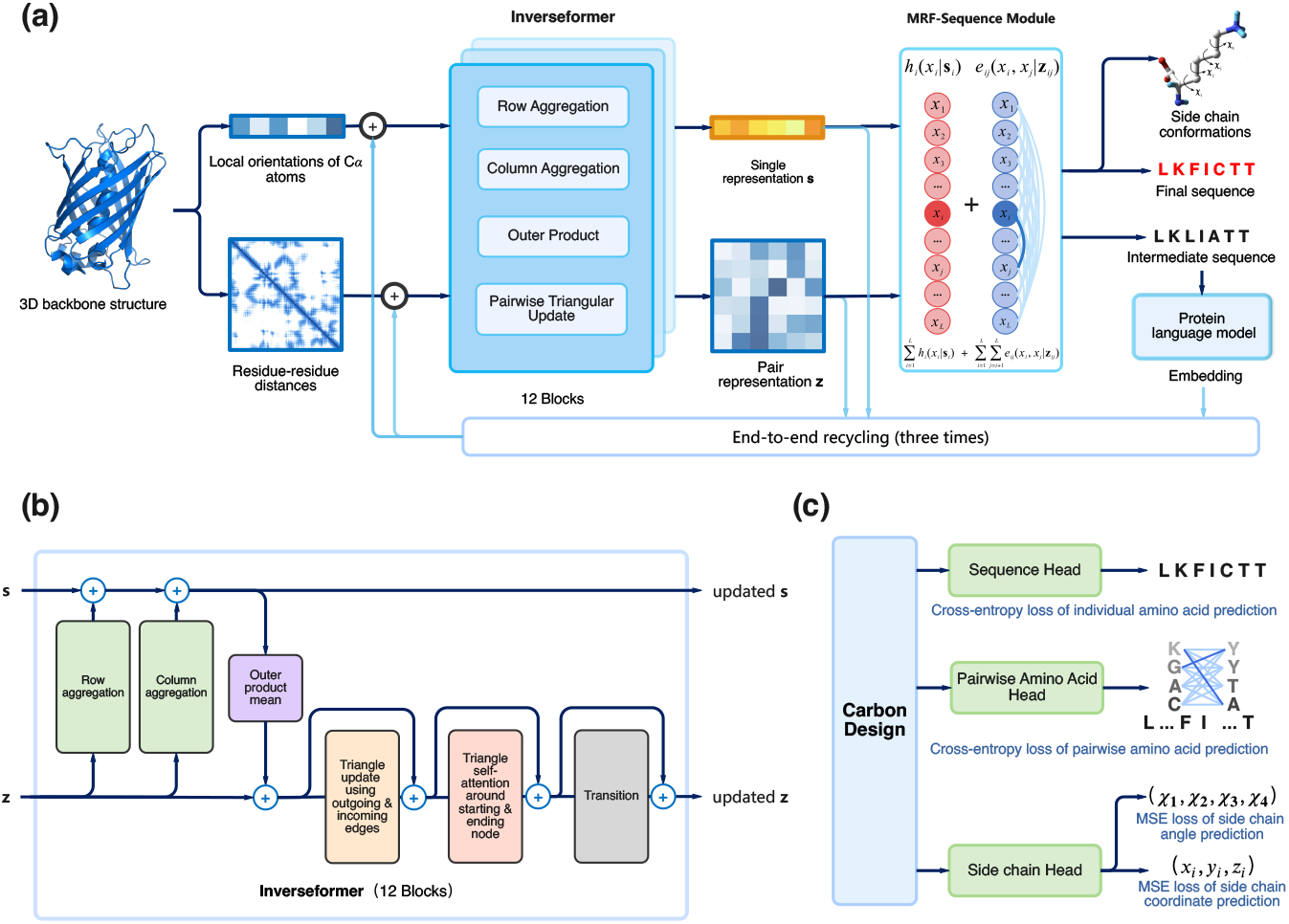
CarbonDesign architecture. **a**, The arrows illustrate the flow of information in the network, designing a 1D protein sequence from a 3D backbone structure. **b**, The Inverseformer blocks update the single and pair representations through node aggregation and triangular edge update layers. **c**, CarbonDesign employs multi-task learning with various training losses, including single and pair amino acid losses, and losses for side chain structures.

The network comprises two main stages. Firstly, we use an Inverseformer module to progressively update the single and pair representations, which are initialized with local orientations and residue-residue distances. Secondly, we use a Markov Random Fields (MRF)-Sequence Module to decode the sequence, with its pair coupling terms and site bias terms parameterized based on the learned pair and single representations, respectively (see Methods).

Inverseformer aims to learn the single and pair representations from which single-site and pairwise amino acids can be decoded (Figure 1b). Single and pair representations interact and undergo refinement through a series of blocks. Specifically, single representations are updated through row aggregation and column aggregation layers with pair presentations as inputs, enabling information flows from 2D to 1D representations. Subsequently, pairwise representations are revised through an outer product layer and four triangular attention layers.

In protein structure prediction, triangular edge updates are intuitively motivated by the need to satisfy the triangle inequality constraints on residue-residue distances. On the other hand, for sequence design, we establish an intuitive connection between Inverseformer’s triangular updates and the edge message updates in the Belief Propagation (BP) algorithm, which is commonly used for learning and inference in probabilistic graphical models like MRFs and Bayesian networks [23, 24]. In the BP algorithm, node and edge messages are updated alternately to aggregate probability mass from neighboring variable nodes. Each edge message *ij* is updated through a triangular edge updates operation, involving all other edge messages *jk* related to variable node *j* (Supplementary Figure S1). Based on this intuition, we hypothesize that the triangular edge updates encourage representations that generate sequences with higher likelihoods under the MRFs model in the following MRFs-Sequence Module.

MRFs-Sequence Module is to construct a probabilistic model for the sequences conditioned on learned single and pair representations. MRFs are widely utilized in direct coupling analysis to model sequence likelihoods [25– 27]. In the context of CarbonDesign, the learned single and pair representations naturally parameterize the coupling and site bias terms in MRFs. Subsequently, a simple ad-hoc algorithm is used to sample the candidate amino acid sequences from the MRFs model (see Methods).

### End-to-end network recycling with a protein language model

The end-to-end network recycling technique enhances model capacity by stacking and reusing the same model architecture with shared weights. Rather than making direct predictions in a single step, this technique employs a self-correcting mode to progressively refine an initial solution by incorporating feed-backs from error predictions. It has been successfully applied within the field of computer vision [18, 19], as well as in AlphaFold for protein structure prediction.

Network recycling enables the model to extract additional features as error feed-backs from the intermediate predictions. In the case of CarbonDesign, learned single and pair representations from the previous recycling rounds serve as features for the next round.

Furthermore, the recycling technique enables CarbonDesign to leverage evolutionary constraints encoded in protein language models like ESM2 in an end-to-end manner. Specifically, the intermediate sequence is first predicted using the single representations, and its embedding is extracted from the language model ESM2 as additional recycling features. Protein language models have the capability to learn efficient representations from millions of sequences and have been successfully applied in predicting protein functions and structures [20]. In the context of CarbonDesign, the language model serves as a prior for the generated sequences.

### Multi-task learning with sequence design and side chain structure prediction

We employ a cross-entropy loss for individual amino acids and an auxiliary cross-entropy loss for pairwise amino acid identities to directly guide the learning of the single and pair representations, respectively. To approximate the exact likelihood of the sequences in the MRFs model [25], we utilize a composite likelihood during training. Moreover, we incorporate a side chain torsional angle loss and a side chain structure loss in training [16], enabling CarbonDesign to predict both the sequences and the corresponding side chain structures (Figure 1**c**).

### Evaluting CarbonDesign on independent testing sets

We extensively evaluated CarbonDesign on two prominent datasets: the CAMEO test set, consisting of 642 structures used in ongoing CAMEO assessments between February 2022 and February 2023 [21], and the CASP15 test set, comprising 65 publicly released structures [22]. We compared our approach with representative methods in protein sequence design, including ProteinMPNN [10], ESM-IF [28], ABACUS-R [13], and ProDESIGN-LE [14].

We evaluated the performance of CarbonDesign using two key metrics: sequence recovery rate and the BLOSUM score [29]. The sequence recovery rate assesses the model’s ability to design sequences that closely match the target structure, while the BLOSUM score measures the similarity between the designed sequences and the native sequences.

Compared to the representative methods, CarbonDesign achieves a sequence recovery rate of 60.1%, outperforming ProDESIGN-LE (42.8%), ABACUS-R (43.6%), ProteinMPNN 020 (default model, with 0.02Å noise) (51.3%), ProteinMPNN 020 (with 0.20Å noise) (45.3%), and ESM-IF (54.8%). Similarly, CarbonDesign outperformed these methods by 1.18, 1.01, 0.95, 0.52, and 0.33 in BLOSUM score.

CarbonDesign continued to exhibit superior performance with the CASP15 test set. It acheives a sequence recovery rate of 53.8%, surpassing ProDESIGN-LE (40.2%), ABACUS-R (38.2%), ProteinMPNN 020 (42.0%), ProteinMPNN 002 (48.1%), and ESM-IF (50.4%) (Figure 2**a**). CarbonDesign also achieves a BLOSUM Score of 2.77, outperforming ProDESIGN-LE (1.80), ABACUS-R (1.74), ProteinMPNN 020 (1.96), ProteinMPNN 002 (2.41), and ESM-IF (2.54) (Figure 2**b**). Remarkably, we have observed that utilizing a larger language model, ESM-3B, leads to a further improvement in sequence design accuracy (Figure 2**e**).

**Fig. 2:**
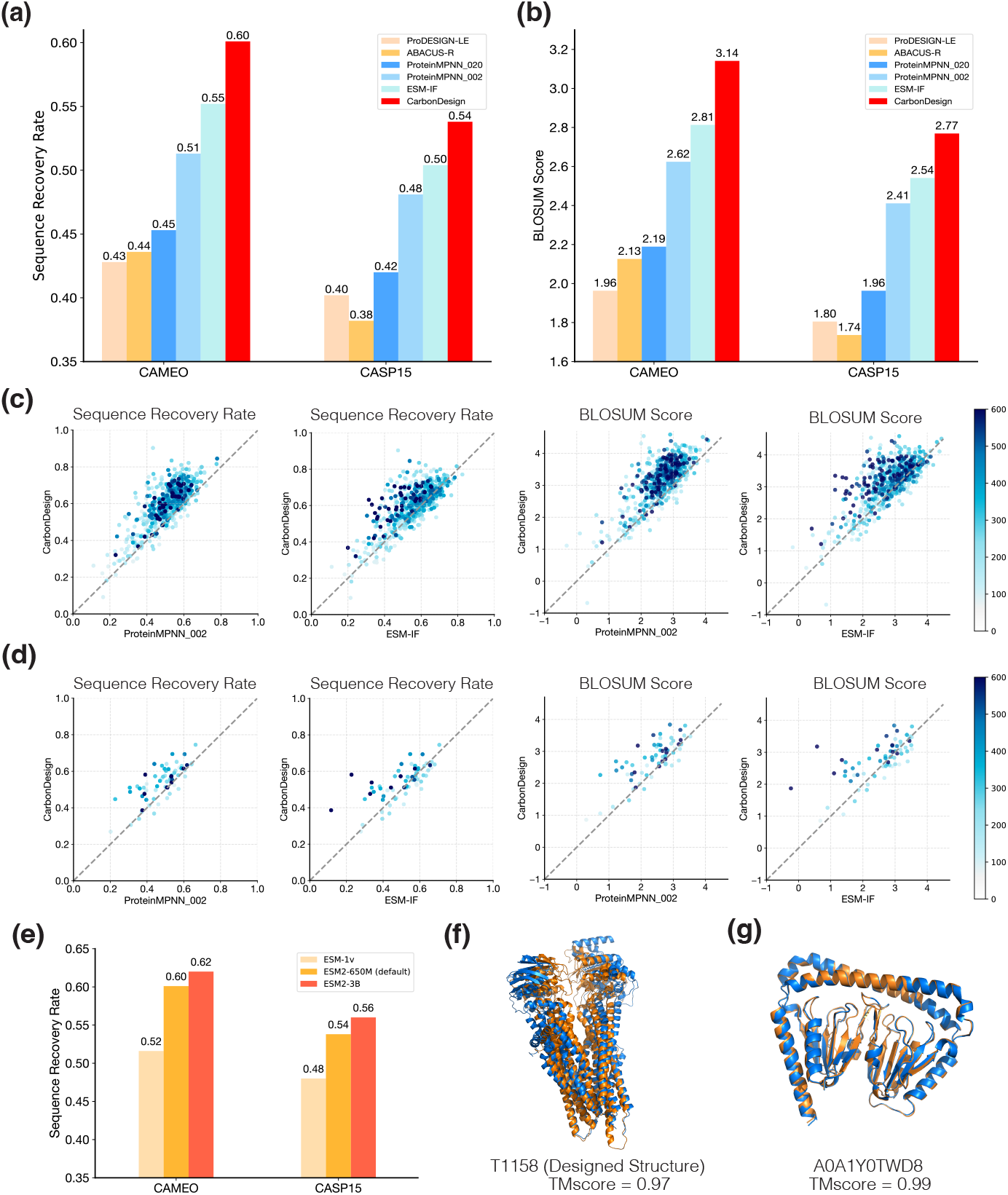
Evaluation of CarbonDesign with the CAMEO and CASP15 independent testing sets. **a-b**, Evaluation with sequence recovery rate and BLOSUM score, respectively. **c-d**, Head-to-head comparisons with other representative methods on CAMEO and CASP15 testing sets, respectively, with color intensity indicating sequence lengths. **e**, Evaluation of CarbonDesign with various protein language models based on sequence recovery rate. **f**, Illustrative case of a long protein T1158 (length: 1340) showing the native structure (blue) and the predicted structure of the designed sequence (orange). **g**, Illustrative case of the novel fold protein dwNTPase mined from AlphaFold DataBase, with the predicted structure of native sequence (blue) and designed sequence (orange).

We further evaluated CarbonDesign using a dataset of orphan proteins characterized by limited or no homologous sequences and a lack of structure templates. These proteins pose a significant challenge for existing structure prediction methods due to the scarcity of evolutionary information [16, 17, 20, 30, 31]. They also serve as a rigorous test set for protein sequence design, as they lack homologous information in existing sequence and structure databases. In our evaluation on the orphan proteins from CASP15, CarbonDesign still demonstrated robust performance, achieving a sequence recovery rate of 49.1% and outperforming all other representative methods (Supplementary Table S6).

Recent advancements in diffusion-based methods have enabled the design of long backbone structures, which pose a challenge for protein sequence design. We curated a dataset of long proteins (*>* 800 amino acids) from CASP15 and CAMEO test sets to evaluate CarbonDesign’s performance. Notably, CarbonDesign achieved a sequence recovery rate of 55.0%, surpassing the compared methods. As an illustrative example, we evaluated CarbonDesign on the multidrug-resistant protein T1158 (Bos taurus MRP4) with a length of 1340 amino acids (Figure 2**f**) (Supplementary Table S5). CarbonDesign demonstrated a sequence recovery rate of 58.1% and a TM-score of 0.97 when comparing the predicted structure via ESMFold with the native structure.

As a case study, we examine the protein Dual-wield NTPase (dwNTPase) (Figure 2**g**) [32], which exhibits a highly novel architecture discovered through data mining of predicted structures in the AlphaFold DataBase [33]. CarbonDesign successfully generates a sequence with a high sequence recovery rate of 70.2%. Furthermore, the predicted structures of the native and the designed sequence exhibit a high similarity (TM-score=0.99). This case highlights the robustness of CarbonDesign with predicted backbone structures and its strong model generalization, enabling accurate designs for novel fold types.

### Improving *de novo* protein design with CarbonDesign

Recent diffusion-based methods, such as RFdiffusion, have revolutionized *de novo* protein design by generating novel backbone structures across diverse fold types that have never been observed in nature. In light of these advancements, we evaluate the efficacy of CarbonDesign in enhancing protein *de novo* design by generating more accurate sequences for these novel backbone structures.

Since native sequences are unavailable for evaluating sequence recovery rate and BLOSUM similarity score, we employ the self-consistency TM-score (scTM) as an alternative measure. Specifically, we first utilize ESMFold to predict the structures of the designed sequences corresponding to the backbone structures generated by RFdiffusion. We then use TM-score to measure the consistency between predicted and original structures. We also note that while scTM is commonly used as a surrogate when native sequences and crystal structures are unavailable, its reliability is contingent upon the accuracy of protein structure prediction.

Following ProteinMPNN and ESM-IF, we introduced noise into the crystal structures during training. This approach accounts for the fact that in practical applications, *de novo*-generated structures or predicted structures may not exhibit the same level of precision as crystal structures commonly used in training. We generated 2560 backbone structures of variable lengths (ranging from 200 to 600) using RFdiffusion and evaluated the performance of CarbonDesign and ProteinMPNN with different noise levels.

Our results highlight two main findings. Firstly, ProteinMPNN consistently outperforms ProteinMPNN in terms of scTM at each noise level (Figure 3**b**). Secondly, we observed that more significant noise levels improve the performance of both CarbonDesign and ProteinMPNN, indicating the beneficial role of noise in generating sequences for *de novo* structures. More specifically, CarbonDesign demonstrates superior performance over the existing representative methods, including ProteinMPNN and ESM-IF, across all different lengths (Figure 3**a**).

**Fig. 3:**
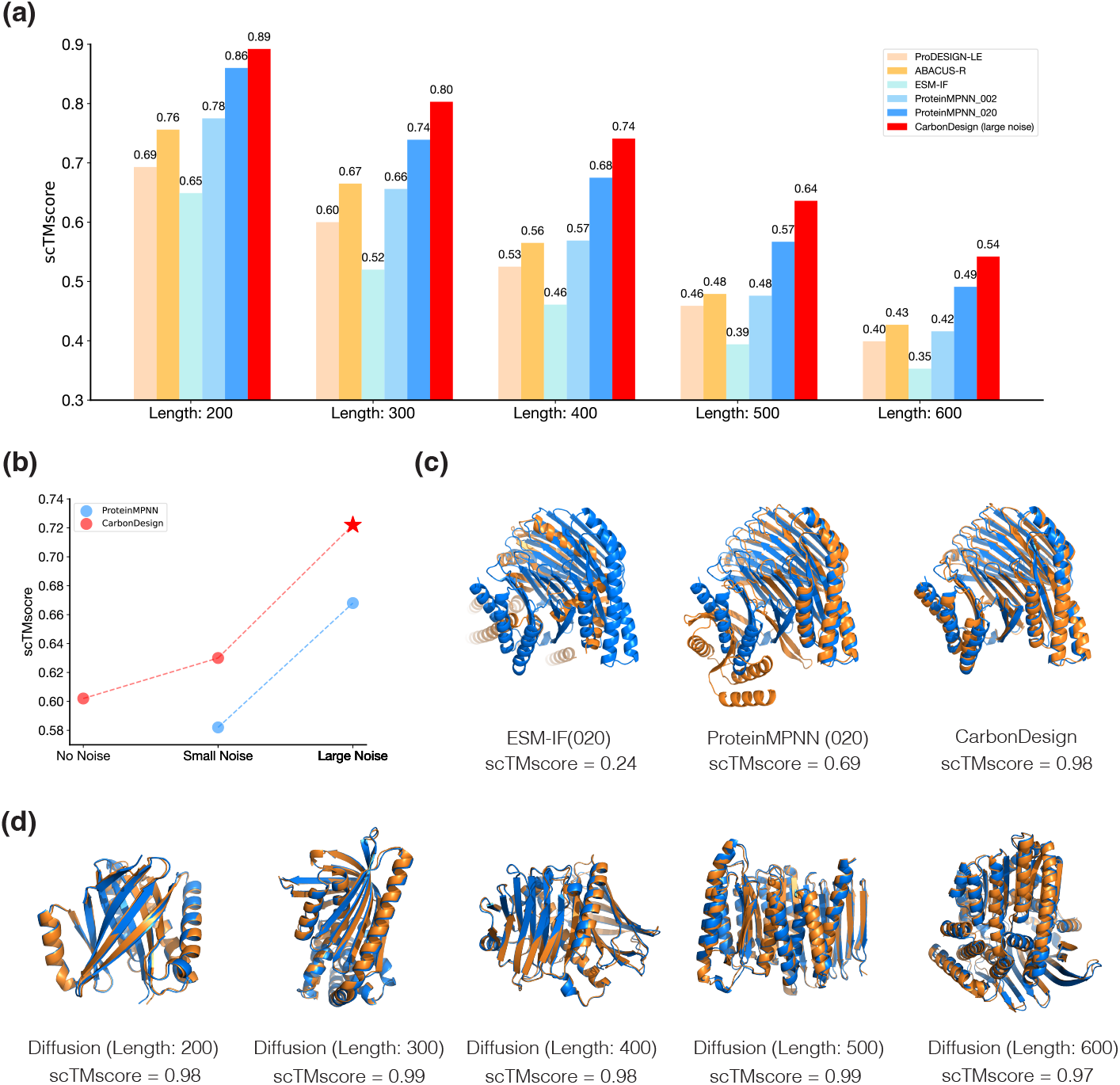
Evaluation of CarbonDesign on *de novo* backbone structures from RFdiffusion. **a**, Evaluation on backbone structures with varying lengths, measured by scTM score. **b**, Impacts of training noise levels on the performance of CarbonDesign and ProteinMPNN. **c**, Illustrative case showing a *de novo* backbone structure (blue) and predicted structures (orange) of designed sequences from ESM-IF, ProteinMPNN, and CarbonDesign, respectively. **d**, Additional illustrative cases of *de novo* backbone structures (blue) with varying lengths and predicted structures (orange) of designed sequences from CarbonDesign.

To assess the broad applicability of CarbonDesign in enhancing protein *de novo* design, we extend our evaluation to include FrameDiff, another recent diffusion-based method. By replacing ProteinMPNN with CarbonDesign, we observe a substantial 0.083 improvement in the scTM score (Supplementary Figure S3), demonstrating the efficacy of CarbonDesign in enhancing the performance of FrameDiff.

Moreover, we present a successful example of a generated backbone structure consisting of 500 residues. CarbonDesign achieves an scTM of 0.98, which is significantly higher than ESM-IF (scTM=0.24) and ProteinMPNN (scTM=0.69) (Figure 3**c**). Furthermore, we demonstrate other successful examples of designed sequences of variable lengths (Figure 3**d**).

### Predicting functional effects of variants via CarbonDesign

The accurate interpretation of the functional effects of variants is crucial in directed evolution-based protein engineering [34, 35], as well as in the context of human genetic studies and clinical testing [36, 37]. Pre-trained language models have emerged as effective zero-shot predictors, alleviating the issue of limited labeled data and mitigating potential human biases in variant annotation [38]. We now show that CarbonDesign also supports zero-shot learning for functional effects prediction, indicating its ability to capture the inherent sequence-structure-function relations.

We first use AlphaFold to predict the protein structures for the testing sequences, which serve as inputs of CarbonDesign. Subsequently, to score the mutational effects of variants on a particular sequence, we calculate the ratio between the likelihoods of the mutated and wild-type sequences based on the CarbonDesign model (see Methods).

We evaluate CarbonDesign on deep mutational scanning datasets with experimentally determined functional scores [39]. CarbonDesign achieves a Spearman correlation of 0.43, outperforming the purely language model-based approaches like ESM-1v and ProGen2 (Figure 4**a**). Furthermore, integrating the scores of CarbonDesign and the other two methods improves the performance, resulting in a Spearman correlation of 0.47. This highlights that CarbonDesign, as a structure-based method, can improve the interpretation of functional effects in combination with purely language model-based methods. We next assess CarbonDesign in predicting the pathogenicity of human genetic variants. Specifically, we focus on four well-known disease risk genes (*BRCA1* [40], *TP53* [41], *PTEN* [42], and *MSH2* [43]) that have a substantial number of high-quality clinical labels in ClinVar. CarbonDesign demonstrates excellent predictive capability for these genes, achieving an average auROC of 0.93 (Figure 4**b**). Notably, CarbonDesign achieves near-perfect separation of benign and pathogenic variants for *TP53* and *PTEN*, with auROC values exceeding 0.95. Furthermore, our results indicate that CarbonDesign out-performs pure language model-based approaches on average in this context (Supplementary S7).

**Fig. 4:**
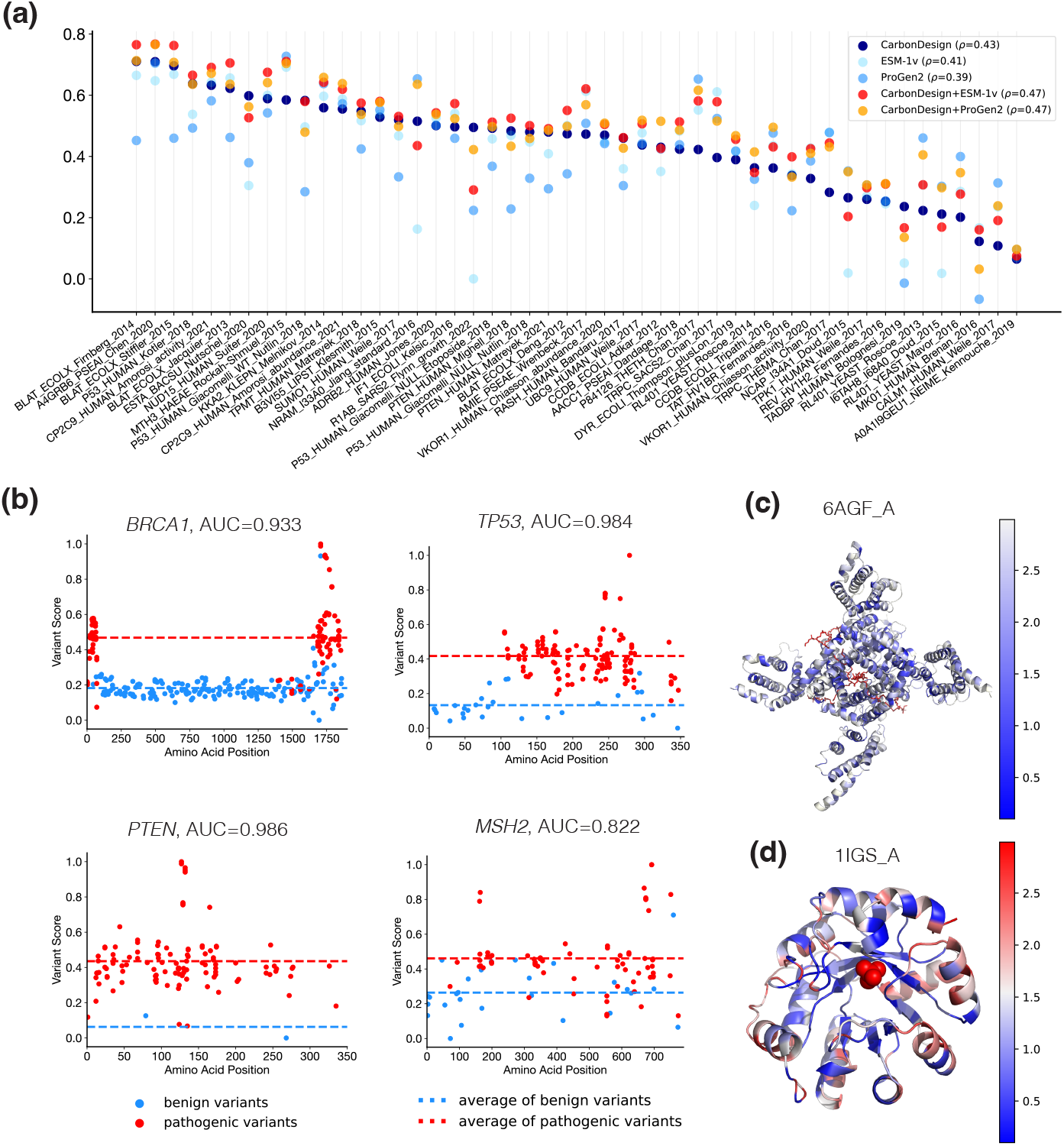
Evaluation of CarbonDesgin in interpreting functional effects of variants. **a**, Evaluation on variants from 49 deep mutational scanning essays. The *x*-axis represents the names of the proteins in the essays, and the *y*-axis represents the Spearman correlation coefficient. **b**, Evaluation on clinical labeled variants in ClinVar for four well-known disease risk genes: *BRCA1, TP53, PTEN, MSH2*. The *x*-axis represents the positions of variants on the proteins, and the *y*-axis represents the functional scores predicted by CarbonDesign. **c**, Entropy variation of protein *Nav*1.4 −*β*_1_, with each position, color-coded based on the level of entropy. Blue regions indicate areas of low entropy, white regions indicate areas of high entropy, and red indicates other binding peptides. **d**, Entropy variation of protein *indole-3-glycerol phosphate synthase*, with each position, color-coded based on the level of entropy. Blue indicates areas of low entropy, red indicates areas of high entropy, and red ions represent phosphate ions.

Furthermore, we observed a correlation between the predicted amino acid distribution and the protein structures. We utilize the entropy of the predicted amino acid distribution as a metric of conservation, with lower entropy indicating higher conservation. As a proof of concept, we examine two proteins, *Nav*1.4 −*β*_1_ [44] (Figure 4**c**) and *indole-3-glycerol phosphate synthase* [45] (Figure 4**d**). In both cases, regions with lower entropy coincide with hydrophobic core regions, associated with functional regions such as the sodium channel and phosphate binding sites.

### Interpreting the CarbonDesign

We trained and evaluated several ablation models to evaluate the relative contributions of the key architecture to CarbonDesign accuracy. These studies focused on examining the impact of the side chain prediction head, end-to-end network recycling technique, and the auxiliary pair head in the MRF model.

CarbonDesign utilizes the side chain head to generate side chain structures of all possible amino acids at each position. We evaluated the prediction accuracy of side chains using the CAMEO and CASP15 datasets and investigated the contribution of side chain heads for sequence design accuracy.

CarbonDesign achieves an average Root Mean Squared Distance (RMSD) of 0.805 (Figure 5**a**). Moreover, the side chain prediction accuracy strongly correlates with the structural context constraints, measured by the number of *C*_*β*_ atoms within an 8Å radius around each residue. Higher side chain prediction accuracy was observed for more constrained residues. For instance, the side chain head of CarbonDesign demonstrated higher prediction accuracy with an RMSD of 0.683 for the protein T1159 (PDB ID: 7PTZ [46]) (Figure 5**c**).

**Fig. 5:**
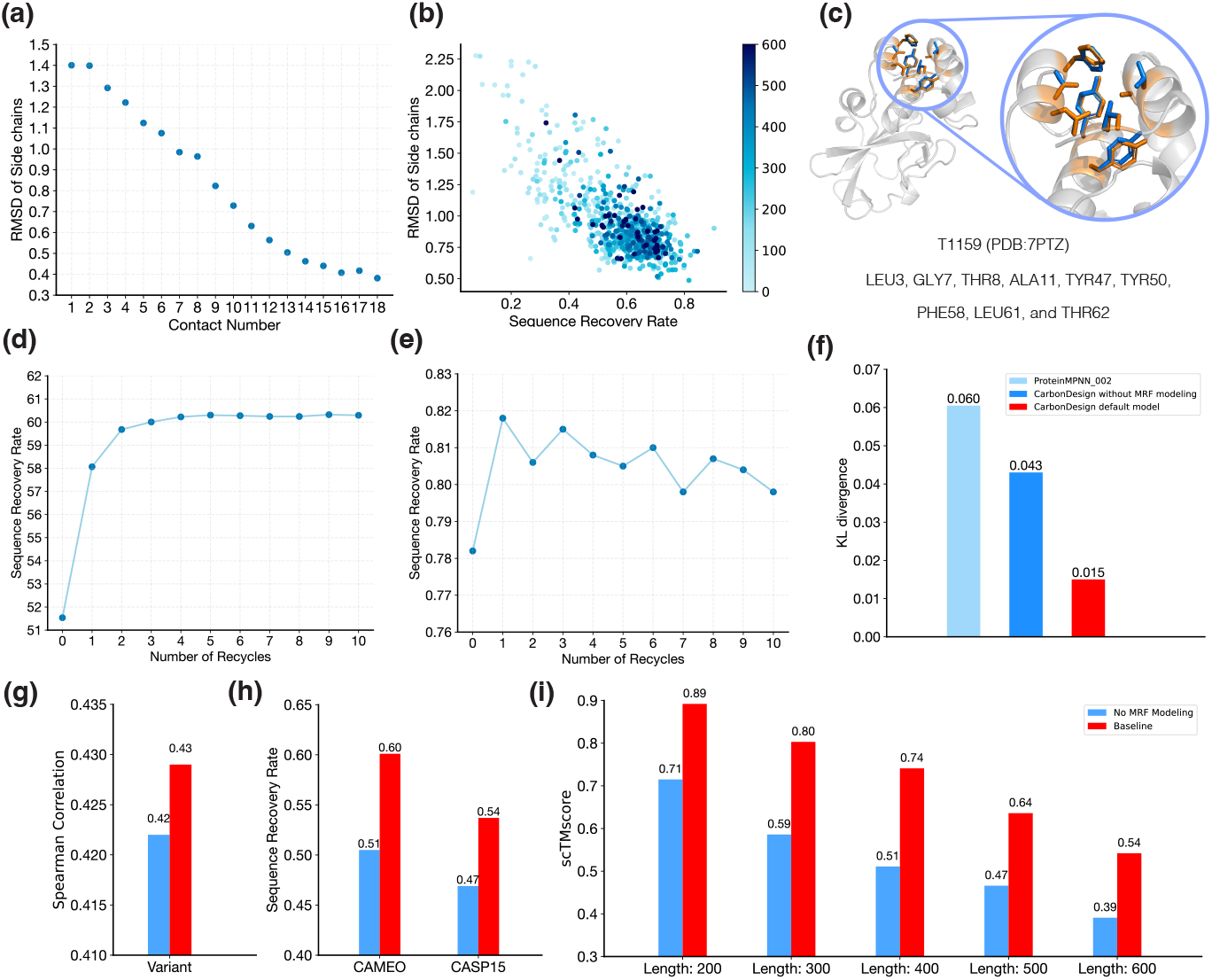
Evaluation of ablation models of CarbonDesign. **a**, Correlation between RMSD error of side chain structure prediction and the number of *C*_*β*_ atoms within an 8Å radius around each residue. **b**, Correlation between sequence design accuracy and side chain structure prediction accuracy on CAMEO and CASP15 datasets. The *x*-axis represents the sequence recovery rate, and the *y*-axis represents the RMSD between predicted and native side-chain structures. **c**, Illustrative case of protein T1159 with predicted side chain structures. Positions of LEU3, GLY7, THR8, ALA11, TYR47, TYR50, PHE58, LEU61, and THR62 are shown, with predicted structures in orange and native structures in blue. **d**, Evaluation of CarbonDesign with varying recycling times, measured by sequence recovery rate on CAMEO and CASP15 testing sets. **e**, Evaluation of CarbonDesign with varying recycling times, measured by scTM score on the backbone structures from RFdiffusion. **f**, KL divergence of the amino acid distribution between designed sequences and the sequences from CAMEO and CASP15 datasets. **g-i**, Evaluation of the effects of pair head in MRFs modeling on performance in deep mutational scanning testing set, CAMEO and CASP15 testing sets, and *de novo* backbone structures from RFdiffusion, respectively. The blue represents the default CarbonDesign model, and the red represents the model with the pair head in MRFs model excluded.

There also exhibits a strong correlation between the side chain prediction accuracy and sequence design accuracy, with a Pearson correlation of 0.73 (Figure 5**b**). The more constrained structural context leads to improved prediction accuracy for both side chain prediction and sequence design tasks, consistent with prior studies [8, 10]. Additionally, training a modified model with the side chain head removed demonstrates the beneficial effect of the side chain head in enhancing the accuracy of designed sequences (Supplementary Table S1).

Network recycling allows the model to incorporate the protein language model in an end-to-end manner. We further assess the contribution of network recycling and the additional sequence embedding from the language models during the recycling stages. Increasing the number of recycling iterations results in an improved sequence recovery rate of designed sequences (Figure 5**d**). We note that when the number of recycling iterations is set to zero, the sequence embedding from the protein language model is not utilized, and the increase in prediction accuracy from no recycling to just one recycling iteration demonstrates the contribution of the protein language model to the predictions. Additionally, network recycling and the protein language model enhance *de novo* protein design evaluated on the backbone structures from the diffusion generative model (Figure 5**e**).

We next explore the accuracy of sequence design at protein core and surface regions. CarbonDesign demonstrates notably higher accuracy at core regions compared to surface regions (Supplementary Figure S4), in line with prior research [10]. Furthermore, we investigate the effects of recycling and the language model on distinct protein regions. We observe a higher relative improvement at surface regions (19.1%) than at core regions (10.3%) (Supplementary Figure S4). These results suggest that the evolutionary constraints embedded in the language model can effectively complement the structural constraints, particularly when the structural context is flexibly constrained.

The pair amino acid head in CarbonDesign directly guides the learning of pair representations in Inverseformer and the pair couplings term in the MRF-Sequence Module. We trained a modified model excluding the pair head to evaluate its contribution. Notably, the pair head significantly improves performance for both crystal structures (Figure 5**h**) and *de novo* structures (Figure 5**i**). Furthermore, we investigated the differences between the amino acid distribution in the designed sequences and the native sequences, measured as Kullback-Leibler (KL) divergence. The model with the pair head can generate sequences with a closer amino acid distribution to the native sequences (Figure 5**f**, Supplementary Figure S5). We also observed a slight improvement in predicting the functional effects of the variants with the DMS testing dataset (Figure 5**g**). These findings underscore the efficacy of the pair head in CarbonDesign.

## Discussion

We present CarbonDesign, a novel approach for protein sequence design that incorporates key concepts from recent successful methods in protein structure prediction. Specifically, CarbonDesign utilizes the Inverseformer architecture, network recycling technique, and multi-task learning strategy to enhance sequence design. Our results demonstrate that CarbonDesign outperforms existing methods in generating candidate sequences for crystal structures, predicted structures, and *de novo* structures derived from diffusion generative models, showing its utility in the *de novo* protein design scenario. Moreover, CarbonDesign supports zero-shot learning for predicting the functional effects of sequence variants, highlighting its ability to capture the intrinsic relationships between protein sequences and their functions.

Notably, CarbonDesign can leverage large-scale pre-trained protein language models, improving sequence design performance. Several previous studies have also demonstrated the utility of language models in various computational protein design scenarios. For instance, ProGen2 employs a generative pre-trained transformer (GPT) model to generate sequences with control tags specifying protein properties [47]. Hie et al. utilize general protein language models to efficiently evolve human antibodies, leading to a substantial improvement in antibody binding affinity [48]. Our CarbonDesign uses the network recycling technique for integrating language models in an end-to-end manner into structure-based protein design.

While CarbonDesign primarily focuses on generating sequences for single chains, it can be readily extended to address a broader range of sequence design tasks, such as designs for hetero multimers, oligo multimers, and target-binding proteins. To achieve this, two key adaptations can be implemented. First, the Inverseformer can be modified to incorporate chain identities as input features, enabling the learning of chain-aware representations. This approach aligns with the adaptation made in AlphaFold-multimer for protein complex structure prediction [49]. Second, by managing the decoding order and constraints, MRF-Sequence Module can produce consistent sequences for each chain within the multimeric structure.

In addition to evolutionary and structural constraints that have been encoded in CarbonDesign, the selection coefficient correlating with allele frequency is another information source that is widely used in interpreting variants in human genetics [50–52]. By integrating both sources, we expect CarbonDesign to have the potential for a broader application in identifying risk genes and prioritizing the damaging variants. Further research is needed to explore this potential.

Our work is limited in focusing solely on the *in silico* evaluation of the designed sequences. While *in silico* metrics provide empirical evidence of whether the designed sequences can fold correctly and exhibit the desired function and are commonly used in the existing methods [10, 11, 15], wet-lab experimental validation is crucial for a comprehensive evaluation of CarbonDesign. It could offer valuable insights and opportunities for improvement and remains our main future work.

## Code and Data Availablility

We plan to release our software and the data after the peer-review process is completed.

## Acknowledgments

We would like to thank the National Key Research and Development Program of China (2020YFA0907000), the National Natural Science Foundation of China (32271297, 62072435, 82130055), and the Project of Youth Innovation Promotion Association CAS to H.Z. for providing financial support for this study.

## Author Contributions

H.Z. and D.B. led the project; H.Z. conceived the ideas and implemented the CarbonDesign model and algorithms; M.R. and H.Z. designed the experiments, and M.R. conducted the main experiments and analysis; M.R. wrote the manuscript, H.Z., D.B, and C.Y. revised the manuscript.

## Methods

### Evaluation data sets

#### CAMEO testing set

We compiled a test set of 728 proteins from the recent CAMEO campaign (between 2022-02 to 2023-02). After excluding short proteins with fewer than 80 amino acids, the final test set consisted of 642 proteins.

#### CASP15 testing set

We included all available proteins from CASP15 that were not canceled and then excluded proteins with lengths less than 80, resulting in a test set of 65 proteins (Supplementary Table S2).

#### Testing set of long proteins

To benchmark the performance of long proteins, we collected proteins with more than 800 amino acids from the CAMEO and CASP15 testing sets. We then used MMseqs to filter overlaps between the two sets with a sequence identity of 40% and selected the representative protein from each cluster. The final test set comprises 13 proteins, with an average length of 1239 (Supplementary S3).

#### Testing set of orphan proteins

We curated a testing set of 9 orphan proteins from the CASP15 set (Supplementary Table S6). Following the criteria of orphan proteins in previous work [20], we first perform the standard AlphFold MSA search process against UniRef [53], MGnify [54], and BFD [55] databases using HHBlits [56] and jackhmmer [57]. Our selection was ultimately narrowed down to proteins that have fewer than 100 homologous sequences and failed to produce a template with a TM-score surpassing 0.5. (Supplementary Table S4) *De novo backbone structures*. We employ RFdiffusion to generate *de novo* backbone structures with variable lengths (200, 300, 400, 500, and 600), producing 512 structures for each length. We also utilize FrameDiff to generate another set of *de novo* backbone structures.

#### Deep mutational scanning dataset

To evaluate CarbonDesign’s efficacy in predicting the functional effects of variants, we compiled the experimentally validated variants from deep mutational scanning (DMS) essays. For the proteins lacking solved crystal structures or with incomplete structures, we use AlphaFold to predict their structures as inputs for CarbonDesign. Due to the limited prediction accuracy of AlaphaFold and other prediction methods for long protein sequences, and the substantial computational resources required, we restrict our analysis to proteins with fewer than 600 amino acids from the ProteinGym DMS dataset. The final testing dataset consists of 179023 variants on 49 genes.

#### Genetic variants on disease genes

To access CarbonDesign’s performance in prioritizing human disease-related variants, we collected the clinically labeled variants from the ClinVar database for four well-studied disease risk genes: *TP53, PTEN, BRCA*, and *MSH2*. Each variant in this dataset is annotated as either pathogenic or benign. This data includes 118 pathogenic (positives) and 175 benign (negatives) variants for *BRCA1*, 111 positives and 2 negatives for *PTEN*, 130 positives and 33 negatives for *TP53*, and 69 positives and 31 negatives for *MSH2*, respectively.

### Training data set

We trained CarbonDesign on protein chains in the Protein Data Bank (PDB) released before Jan. 1st, 2020, determined by X-ray crystallography or cryoEM. We only include the structures with a resolution better than 5.0Å and with more than 50 amino acids. Sequences were clustered at 40% sequence identity cutoff using MMseqs2, resulting in 30,828 clusters.

### Input features

CarbonDesgin incorporates inter-residue distances as edge features and local orientations of 4 consecutive *C*_*α*_ atoms as node features.

#### Edge features

Following ProteinMPNN [10], we calculate the distances between *N, C*_*α*_, *C, O* atoms, and virtual *C*_*β*_ atoms for each residue pair. We then divide the distances from 0Å to 15Å into 20 bins. The bin indices are then one-hot encoded and mapped through a feed-forward layer to initialize the pair representations. We note that we mask all the edges whose distance exceeds 15Å. Additionally, following AlphaFold, we incorporate relative positional encoding for edge features.

#### Node features

For each residue at position *i*, we employ the Gram-Schmidt process to calculate the local frame defined by the 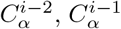, and 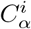 atoms. In this frame, 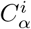 serves as the origin, the direction of 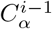 as the *x*-axis, and 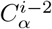 determines the *x*-*y* plane. Specifically, its basis [***a, b, c***] is obtained as follows:

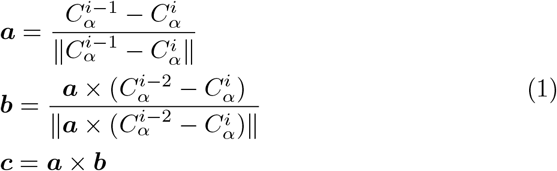

Subsequently, the orientation of 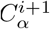 is represented using its local coordinate with respect to this frame (Supplementary Figure S2). Similarly, we calculate the local orientation of 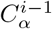 with respect to the 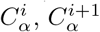, and 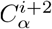 atoms.

### Inverseformer achitecture

We utilize a series of Inverseformer blocks to learn representations from the input backbone structures (Algorithm 1). Each block has a single representation **s**_*i*_ of nodes and a pair representation **z**_*ij*_ of edges as its input and output and processes them through several layers.

We leverage row and column aggregation layers to update the single representations from the pair representations (Equation 2). We note that the aggregation layers are specifically tailored to incorporate edge information directly. The original row and column attention layers in the AlaphaFold Evformer architecture are unsuitable for our purpose, as they primarily focus on aggregating information on nodes, with only a bias on edges.

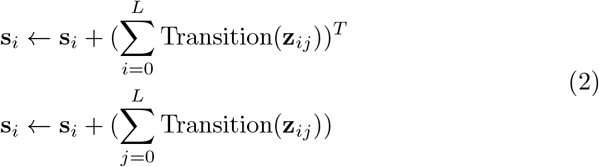

We adopt a similar approach as AlphaFold for updating pair representations. We use an “Outer product mean” block to integrate the single representations, followed by triangular update blocks. Furthermore, we introduce residual connections and dropout layers to prevent overfitting.

The final Inverseformer block produces a highly processed single representation **s**_*i*_ for individual residues and a pair representation **z**_*ij*_ for residue-residue pairs, which contain the necessary information for the MRF-sequence module to decode the sequences. These representations are crucial for accurately predicting the protein sequences.

#### Algorithm 1

Inverseformer

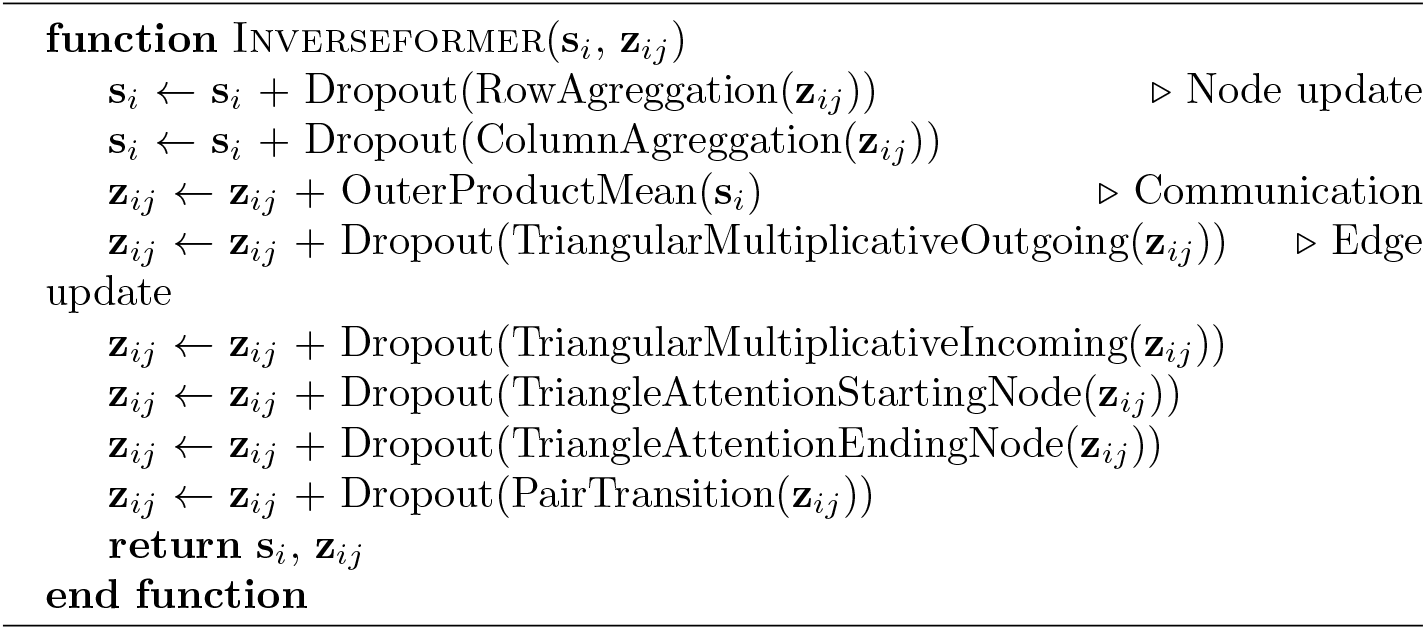

### MRF-Sequence Modeule

We employ an MRF(Markov Random Field)-Sequence Module to decode the sequence from the learned representations. We denote a protein sequence of length *L* as ***x*** and the type of the *i*-th amino acid as *x*_*i*_. And we use the random variable **X** to denote the predicted amino acid sequence.

MRFs have proven to be effective in modeling the distributions of sequences within a protein family [26, 27]. In CarbonDesign, we adopt an amortized Markov Random Field (MRF) model to describe the distribution of the designed sequences (Equation 3), which is conditioned on the learned single representations **s** and pair representations **z**:

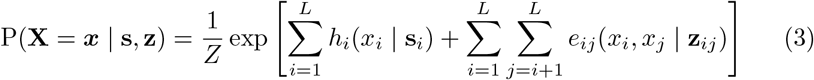

Here, *h*_*i*_ and *e*_*ij*_ are the *conversation bias* term and *pairwise coupling* term, respectively, in the vanilla MRF model, and *Z* is the partition function. For CarbonDesign, we employ a feed-forward layer to project the learned single representation **s**_*i*_ and pair representation **z**_*ij*_ to *h*_*i*_ and *e*_*ij*_, respectively. The training and inference of the MRF model are interconnected with other modules in CarbonDesign and will be elaborated in the subsequent sections.

### Model inference

CarbonDesign consists of two main components: Inverseformer blocks and the MRF-Sequence Module. The Inverseformer blocks take input backbone features as initial representations to compute updated representations. Subsequently, the MRF-Sequence Module utilizes these representations to generate intermediate sequences, final designed sequences, and corresponding side chain structures.

For inference, the whole network is executed sequentially *N*_cycle_ times, where the output single and pair representations of the former execution are recycled as inputs for the next execution (Algorithm 2). During the recycling phase, the intermediate sequence is inferred using the MRF model, and additional recycling features are extracted from the protein language model ESM2 by obtaining embeddings of the sequence.

In the MRF-Sequence Module, we employ both an efficient *local inference mode* and a more accurate *global inference mode* for generating intermediate and final designed sequences, respectively. The *local inference mode* utilizes only the *conservation bias* term to infer the intermediate designed sequence:

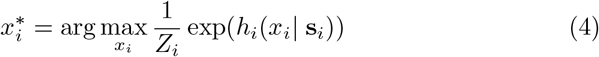

Here, *Z*_*i*_ represents the local partition function involving only the conservation bias terms at position *i*. In contrast, the global inference mode optimizes the sequence by maximizing the sequence probability under the MRF model, considering both the *conservation bias* term and the *pairwise coupling* term (Equation 4). The efficient local inference mode allows obtaining the embeddings of intermediate sequences in a computationally feasible manner. Since exact optimization is challenging for the global mode, we initialize the inference using the sequence from the local inference mode and update sequences using a fast greedy algorithm (Supplementary Note 3).

During the inference stage, when the types of amino acids are unknown, we first utilize the single presentation **s**_*i*_ to predict the side chain structures 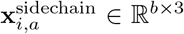 ∈ ℝ ^b×3^for all possible amino acids, where *b* represents the number of side chain atoms and *a* covers 20 amino acid types. The final side chain structures are materialized from 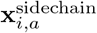 once the final designed sequence is determined by the *global inference mode* of the MRFs model.

### Training losses

The network is trained end-to-end, with gradients coming from the losses for reconstructing native sequences and predicting side chain atomic coordinates. The total per-example loss can be defined as follows:

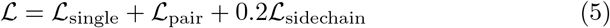

To restore native sequences, we utilize single cross-entropy loss *ℒ*_single_ and pairwise cross-entropy loss *ℒ*_pair_ as direct supervision for the *conservation bias* term *h*_*i*_(*x*_*i*_ | **s**_*i*_) and the *pairwise coupling* term *e*_*ij*_(*x*_*i*_, *x*_*j*_ | **z**_*ij*_), respectively. To calculate *ℒ*_single_, we linearly project the single representations **s**_*i*_ to obtain logits and then compute the cross-entropy loss using the native sequence as labels. For *ℒ*_pair_, we use a pairwise pseudo-likelihood (Equation 6) to approximate the full likelihood of the sequence under the MRF model, following our previous work on residue-residue contacts prediction [25]. For each pair of amino acids in the sequence, its pseudo-likelihood conditioned on other amino acids is given by:

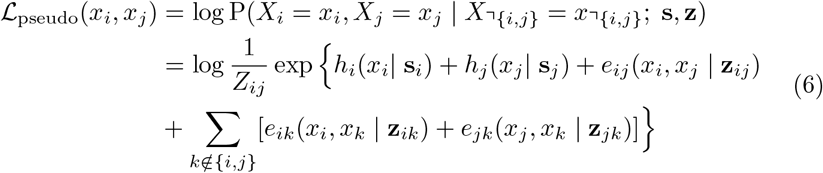

Here, *Z*_*ij*_ is the local partial function. This pseudo-likelihood produces the predicted distribution of amino acid pairs, and *ℒ*_pair_ is computed with pairwise amino acid identities as the labels. We note that *ℒ*_pair_ can directly supervise *e*_*ij*_ in the MRF-Sequence Module and pair representation **z**_*ij*_ in the Inverseformer. Additionally, we added a 0.01 factor of *L*1 and *L*2 regularization terms into *ℒ*_pair_.

The side chain loss consists of three components:

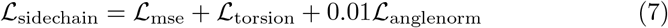

*ℒ*_mse_ is the Mean Squared Error (MSE) for predicted side chain atomic coordinates. Additionally, following AlphaFold, we incorporate the loss terms *ℒ*_torsion_ and *ℒ*_anglenorm_ to evaluate the error of side chain torsion angles [16].

#### Algorithm 2

CarbonDesign Model Inference

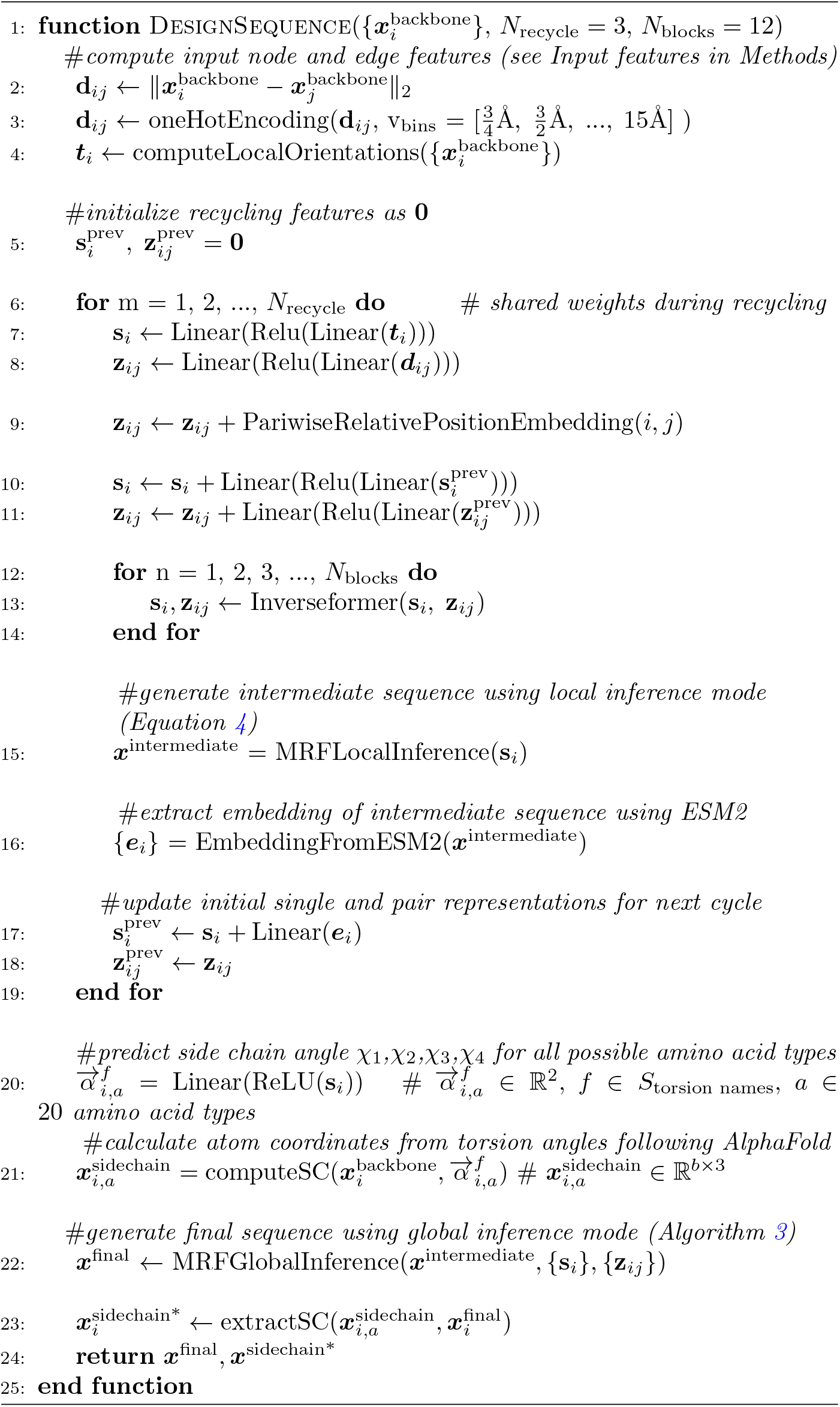

### Additional training details

For training, we utilize the Adam [58] optimizer with a *β*_1_ value of 0.9 and a *β*_2_ value of 0.99. The base learning rate is set to 3e-4 with a warm-up period of 1000 steps, starting from 1e-5, and the training proceeds for an additional 20000 steps. We randomly crop very long proteins during training with a crop size of 400. The network architecture and training pipeline is implemented in PyTorch [59], and training is performed on 16 NVIDIA A40 GPUs.

We trained several ablation models to assess the contributions of different mechanisms utilized in CarbonDesign. Following ProteinMPNN [10] and ESM-IF [15], we add noises to structures during training to deal with noises in *de novo* and predicted backbone structures in practical applications. In the default CarbonDesin model, we added a 0.2Å noise to half of the training samples (referred to as *small noise*). To further investigate the effects of noise levels on the performance with *de novo* backbone structures, we trained two additional models: one without any noise (referred to as *no noise*), and another with a 0.2Å noise applied to all training samples (referred to as *large noise*). For more details on other ablation studies, please refer to Supplementary Table S9.

### CarbonDesign score for predicting functional effects of variants

In CarbonDesign, each variant is scored using the log odds ratio between the mutated and wild-type sequences. The variant score is defined as:

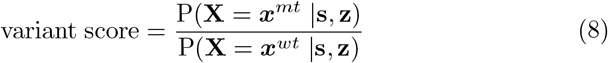

Here, P(**X** = ***x***^*mt*^ |**s, z**) and P(**X** = ***x***^*wt*^ |**s, z**) represents likelihood of the mutated and wild-type sequence, respectively, under the amortized MRFs model (Equation 3).

## 1 Supplementary Materials

### 1.1 Details on running the compared methods

#### ESM-IF

We utlize the test script provided in the ESM GitHub repository (https://github.com/facebookresearch/esm/tree/main/examples/inverse_folding), with the model esm_if1_gvp4_t16_142M_UR50 and all other default settings.

#### ProteinMPNN

ProteinMPNN offers multiple models based on varying noise levels. For a more comprehensive comparison, we use the Protein-MPNN (default) model with 0.2Å noise and the ProteinMPNN (v 48 002) model with 0.02Å noise. We use the testing scripts of ProteinMPNN from the ProteinMPNN GitHub repository (https://github.com/dauparas/ProteinMPNN). Except for our selection of different models for testing, all parameter settings employ the default options provided by GitHub.

#### ProDESIGN-LE

We utilized all sequences designed by the ProDESIGN-LE provided server (http://falcon.ictbda.cn:89/serving2/submit/aFGjrWnGyA/?app=prodesign). All parameters were selected according to the default settings of this method.

#### ABACUS-R

We utilized the test script provided on the GitHub (https://github.com/liuyf020419/ABACUS-R/tree/main/demo) for protein sequence design through ABACUS-R. All parameters were selected from the default options provided in the config file on the website.

#### ESM-1v

In predicting functional effects of variants, we employed ESM-1v as the benchmark criterion. We use the testing script of ESM-1v in the ESM GitHub repository (https://github.com/facebookresearch/esm/tree/main/examples/variant-prediction). All the hyper-parameters are default.

#### ProGen2

We use the model ProGen2-xLarge (6.4B). The GitHub repository is (https://github.com/salesforce/progen/tree/main/progen2). All hyper-parameters are default.

### 1.2 Additional details on ablation models

The ablation models we trained include:

1. Based on CarbonDesign (default), we removed the network recycling, and this model will also disable the language model added during the recycling stages.
2. Based on CarbonDesign (default), we removed the pairwise amino acid head during training.
3. Based on CarbonDesign (default), we removed the side-chain head during training.

**Table S1:**
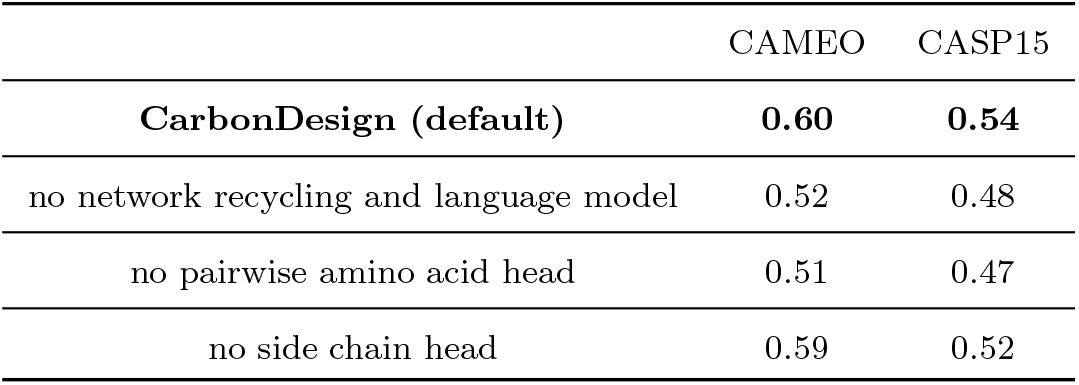
Evaluation of ablation models on CAMEO and CASP15 testing sets.

|  | CAMEO | CASP15 |
| --- | --- | --- |
| **CarbonDesign (default)** | **0.60** | **0.54** |
| no network recycling and language model | 0.52 | 0.48 |
| no pairwise amino acid head | 0.51 | 0.47 |
| no side chain head | 0.59 | 0.52 |

### 1.3 Intuitive connections between Inverseformer and Belief Propagation Algorithm

During the encoding of backbone structures, the **direct** operation on nodes and edges plays a crucial role in determining the information flow and learning of their representations.

ProteinMPNN and ESM-IF utilize different approaches for node and edge encoding. In ProteinMPNN, a graph neural network is used, while ESM-IF employs a Geometric Vector Perceptron (GVP) [60] for this task. Information on each edge in these models is updated based on the edge itself and its related edges (Figure S1**a**).

In contrast, CarbonDesign’s Inverseformer uses triangular attention updates on edges, where the representation of each edge is updated by considering the representations of edges sharing a node (Figure S1**b**). This approach is inspired by AlphaFold’s Evoformer, where triangular edge updates are motivated by the need to satisfy the triangle inequality constraints on residue-residue distances. In CarbonDesign, we establish an intuitive connection between triangular edge updates in sequence design and the Belief Propagation algorithm used in probabilistic graphical models.

In probabilistic models like Bayesian networks and Markov Random Fields, a graph *G* = (*V, E*) is employed to describe the joint distribution of *P* (*X*_1_, *X*_2_, …, *X*_*n*_) for *n* random variables (Figure S1**c**). Each variable *x*_*i*_ is represented as a node, and edges between variables represent direct correlations. The Belief Propagation algorithm aims to calculate the marginal distribution of a specific variable or a subset of variables by iteratively aggregating probability mass from neighboring nodes. Specifically, *m*_*ji*_(*x*_*j*_) represents the “belief” of variable *x*_*i*_ based on variable *x*_*j*_, and it is updated by aggregating information from all edges *jk* (*k≠ i*) connected to node *j* (Equation 9).

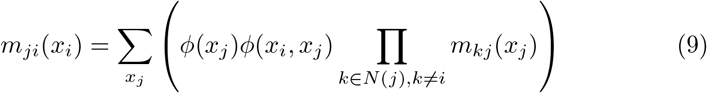

**Fig. S1:**
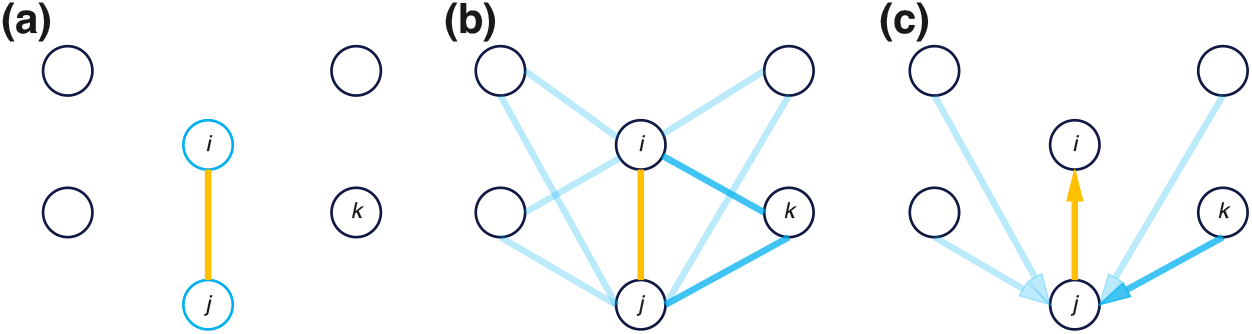
Edge update in ProteinMPNN, Inverseformer, and Belief Propagation algorithm. **a**, ProteinMPNN updates representation of edge *ij* using the edge itself and the nodes *i* and *j*. **b**, Inverserformer updates representation of edge *ij* using the information from all related edges *ik* and *jk* (*i, j≠ k*). **c**, BP algorithm updates the belief on edge *ij* using beliefs from all edges *jk* connected to *j* (*k≠ j*).

### 1.4 Global inference mode for the amortized MRF model

In the MRF-sequence module, we leverage both a *local inference mode* to generate intermediate sequences (see Methods in the main text) and a *global inference mode* to produce the final designed sequences (Algorithm 3). Since it is computationally infeasible to determine the sequences that exactly maximize the full likelihood under the MRF model (Equation 3), we use an efficient and straightforward greedy approach for approximation.

We initialize the sequence with the *local inference mode*, denoted as *x*^*old*^. Subsequently, we update each amino acid by maximizing its conditional likelihood given the identities of other amino acids:

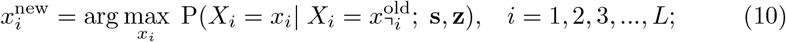

The conditional likelihood involves both the conservation bias term *h*_*i*_(*x*_*i*_ |***s***_*i*_) and the pairwise coupling term *e*_*ij*_(*x*_*i*_, *x*_*j*_ |***z***_*ij*_), and it can be calculated efficiently as follows:

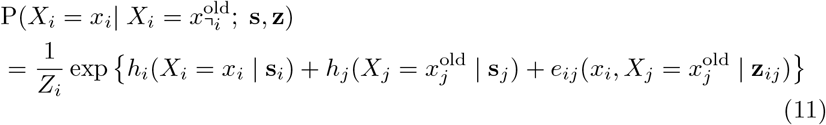

Here, *Z*_*i*_ is the local partition function that sums over all 20 possible amino acid types at position *i*. For both training and inference, we only include edges for neighboring residues within a *C*_*β*_ *−C*_*β*_ distance of 8Å.

We alternately update each amino acid, and after completing updates for the entire sequence, we proceed to the next round of updates until the sequence converges. Typically, sequences converge within 2 rounds of updating, and we set the maximum number of rounds as 3. We note that during the inference of the MRF model, both ***s***_*i*_ and ***z***_*ij*_ are held constant and treated as static inputs, and there is no need to run Inverformer to update them in the process.

#### Algorithm 3

Global inference mode of the MRFs model

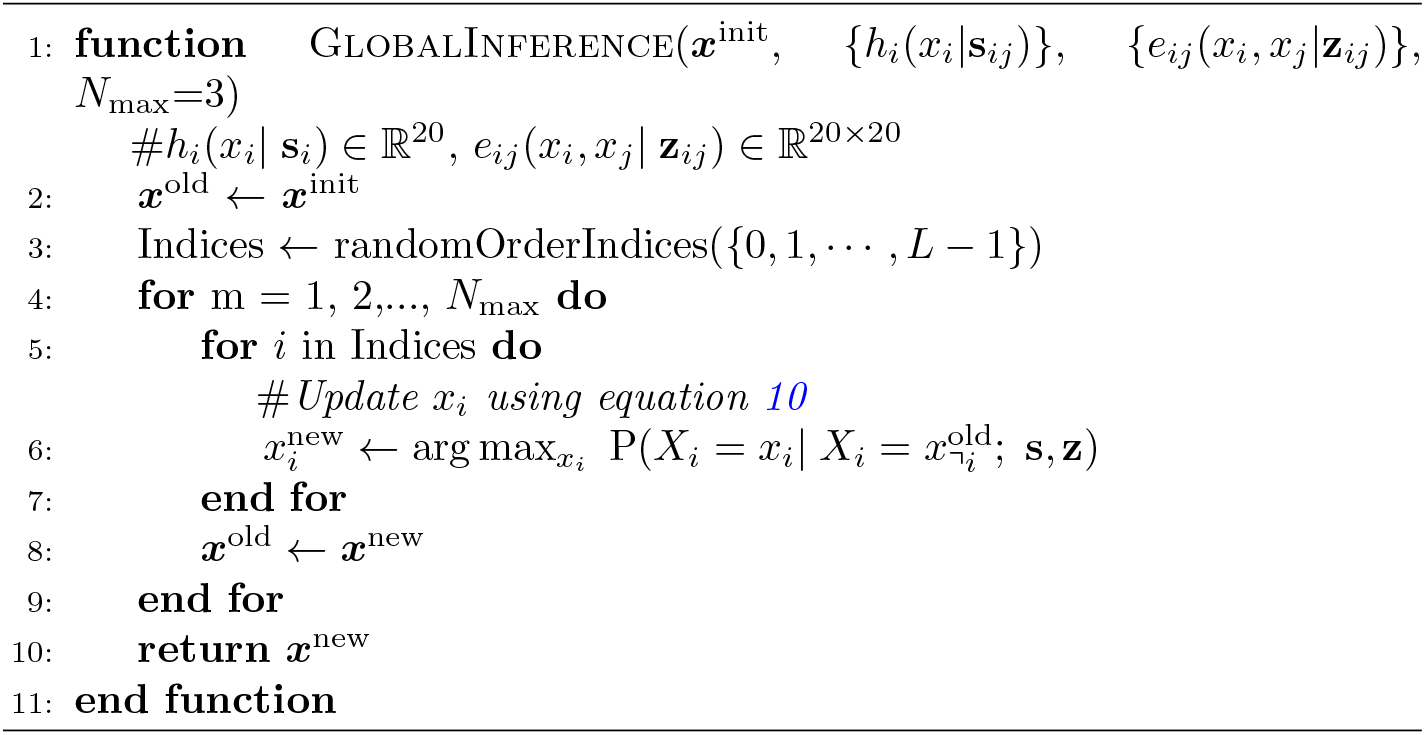

## Supplementary Figures

**Fig. S2:**
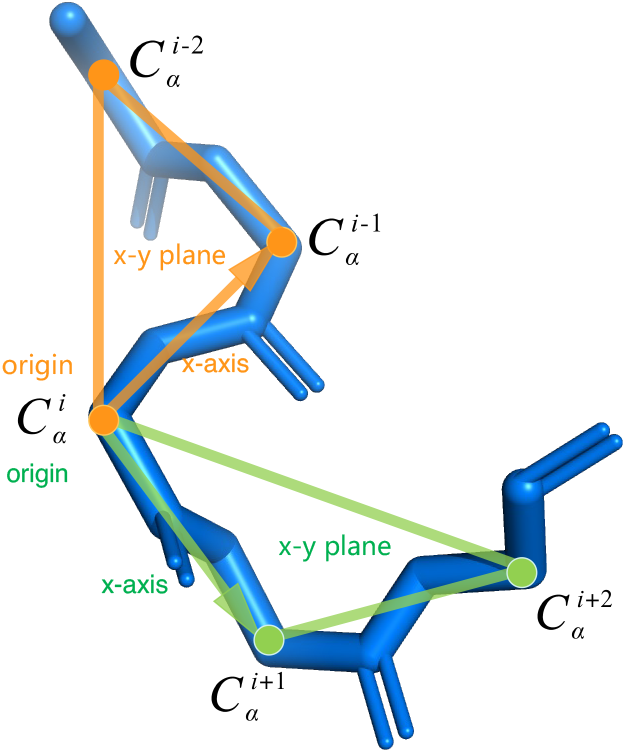
Computing local orientations of *C*_*α*_ atoms in backbone structures. We utilize the Gram-Schmidt process to calculate the local frame formed by the 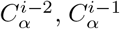, and 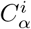 atoms. Subsequently, we represent the local orientation of 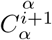 as its local coordinate in the frame. Similarly, we calculate the local orientation of the 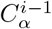 with respect to the 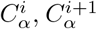, and 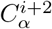 atoms.

**Fig. S3:**
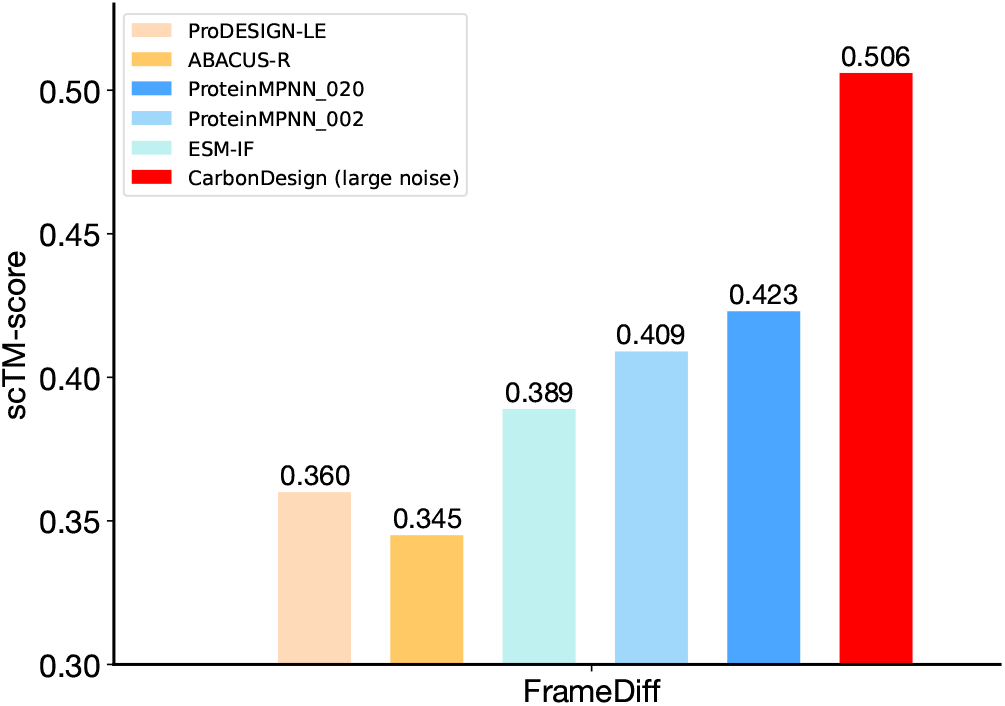
Evaluation of CarbonDesgin on *de novo* backbone structures from FrameDiff.

**Fig. S4:**
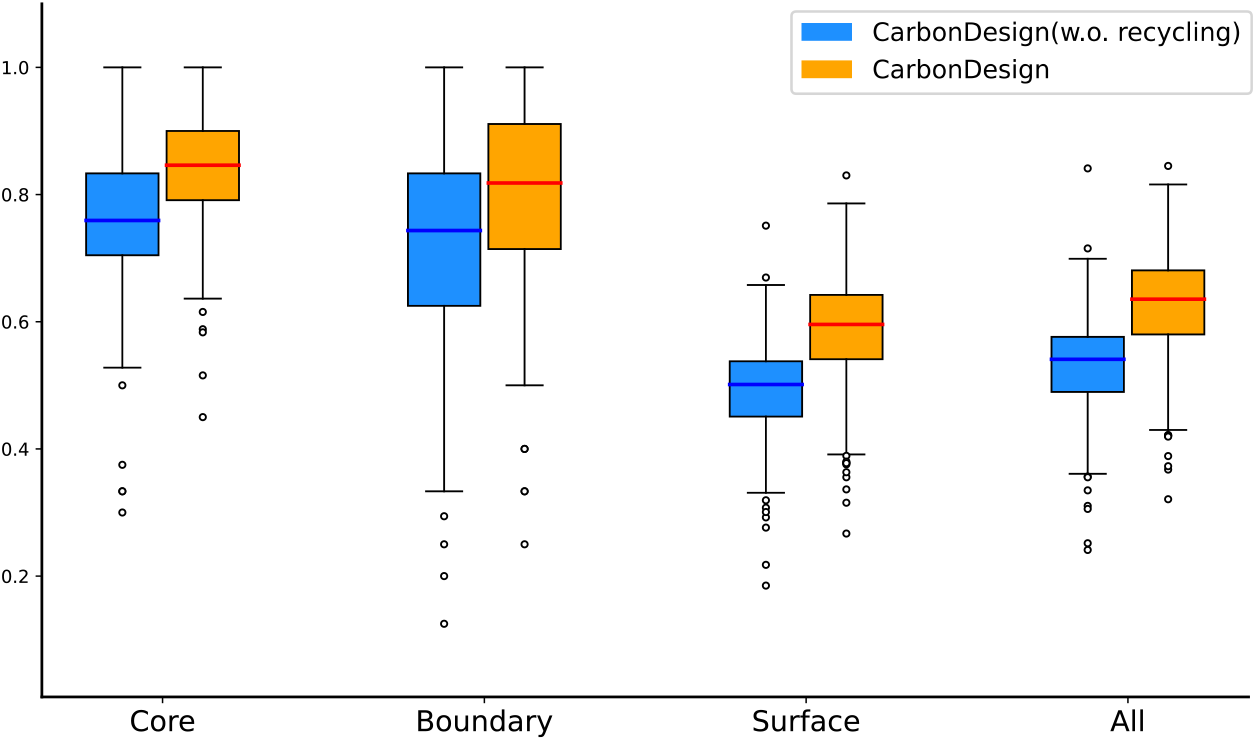
Evaluation of CarbonDesign on protein core and surface regions. The relative solvent-accessible surface area (RSA) for each residue is calculated and categorized into Core (*<* 0.25), Boundary (0.25-0.75), and Surface (*>* 0.75) regions. Sequence recovery rates are evaluated for both the CarbonDesign default model and the model with the recycling and protein language model excluded.

**Fig. S5:**
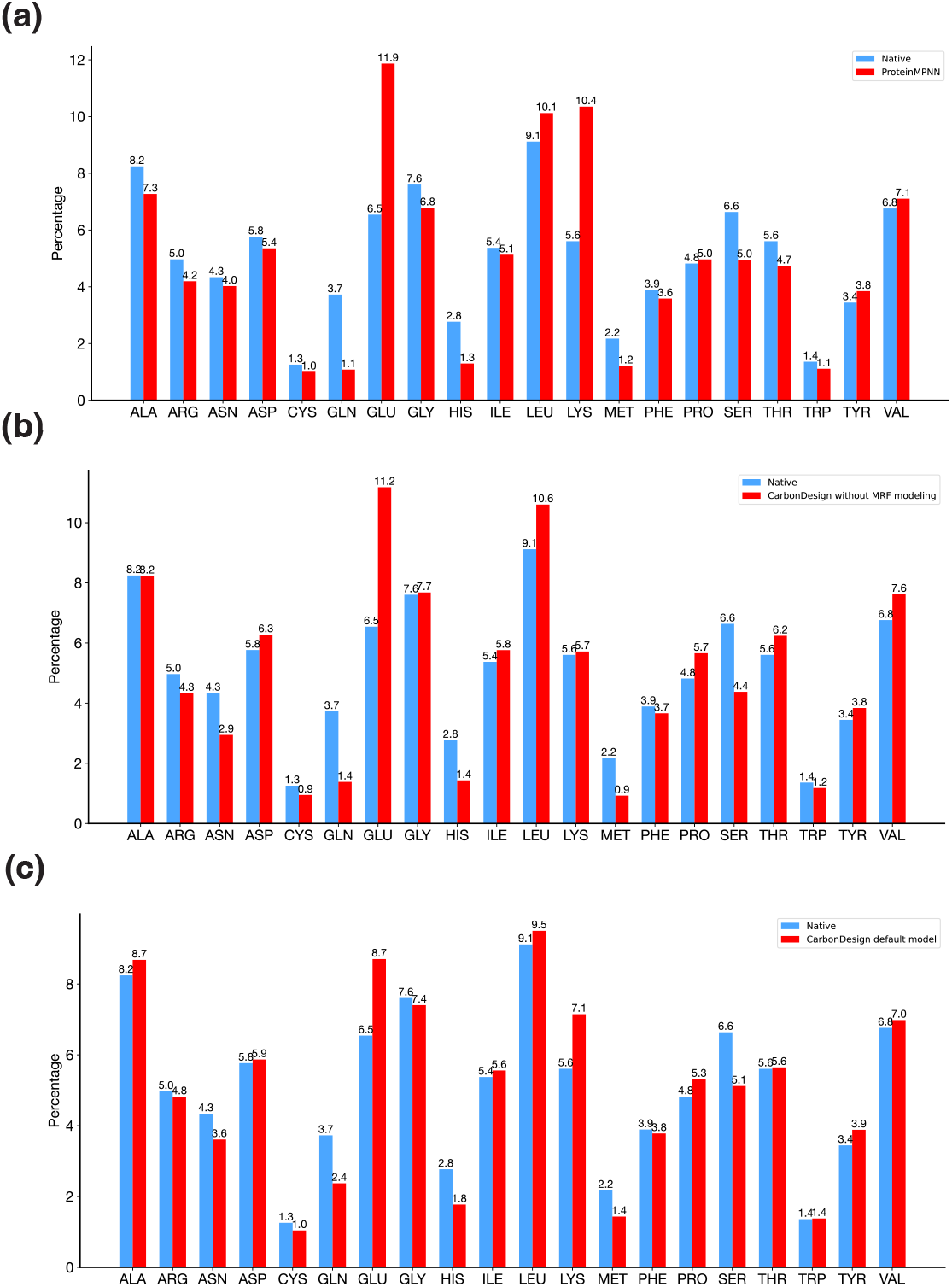
Distributions of amino acid types in designed and native sequences. **a**, A comparison between the distributions of amino acid types in sequences designed by ProteinMPNN and native sequences. **b**, A comparison between the distributions of amino acid types in sequences designed by CarbonDesign without MRF modeling and native sequences. **c**, A comparison between the distributions of amino acid types in sequences designed by CarbonDesign and native sequences.

## Supplementary Tables

**Table S2:**
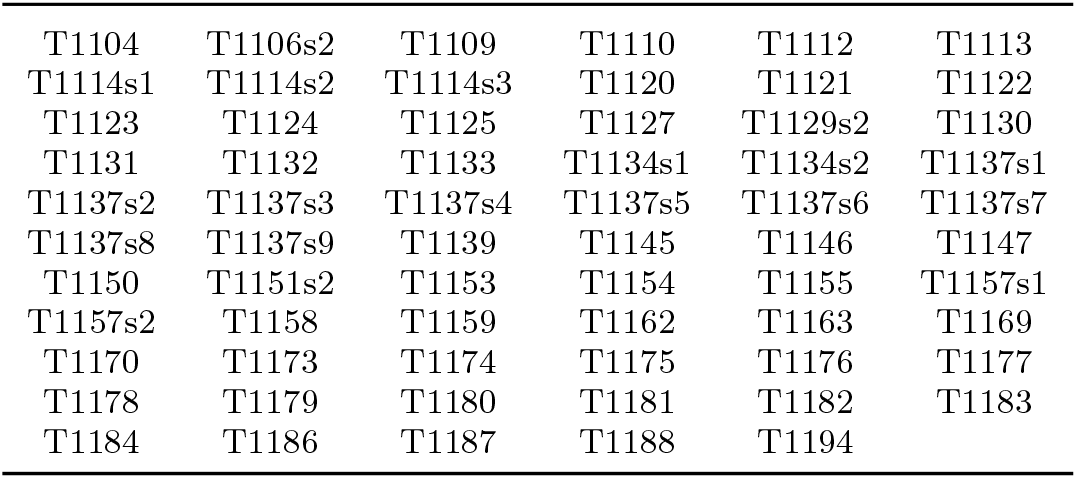
List of protein names in the CASP15 testing set.

|  |  |  |  |  |  |
| --- | --- | --- | --- | --- | --- |
| T1104 | T1106s2 | T1109 | T1110 | T1112 | T1113 |
| T1114s1 | T1114s2 | T1114s3 | T1120 | T1121 | T1122 |
| T1123 | T1124 | T1125 | T1127 | T1129s2 | T1130 |
| T1131 | T1132 | T1133 | T1134s1 | T1134s2 | T1137s1 |
| T1137s2 | T1137s3 | T1137s4 | T1137s5 | T1137s6 | T1137s7 |
| T1137s8 | T1137s9 | T1139 | T1145 | T1146 | T1147 |
| T1150 | T1151s2 | T1153 | T1154 | T1155 | T1157s1 |
| T1157s2 | T1158 | T1159 | T1162 | T1163 | T1169 |
| T1170 | T1173 | T1174 | T1175 | T1176 | T1177 |
| T1178 | T1179 | T1180 | T1181 | T1182 | T1183 |
| T1184 | T1186 | T1187 | T1188 | T1194 |  |

**Table S3:**
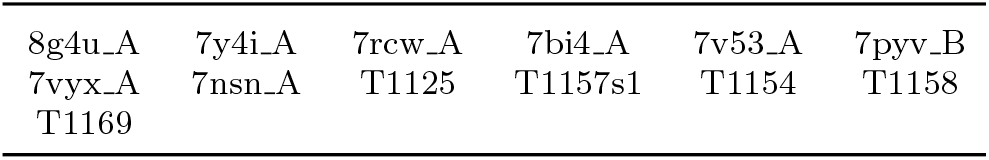
List of protein names in the testing set of long proteins.

|  |  |  |  |  |  |
| --- | --- | --- | --- | --- | --- |
| 8g4u_A | 7y4i_A | 7rcw_A | 7bi4_A | 7v53_A | 7pyv_B |
| 7vyx_A | 7nsn_A | T1125 | T1157s1 | T1154 | T1158 |
| T1169 |  |  |  |  |  |

**Table S4:** List of protein names in the testing set of orphan proteins.

|  |  |  |  |  |  |
| --- | --- | --- | --- | --- | --- |
| T1122 | T1130 | T1131 | T1125 | T1113 | T1178 |
| T1184 | T1155 | T1129s2 |  |  |  |

**Table S5:** Evaluation on the testing set of long proteins measured with sequence recovery rate. The table presents the results for 13 proteins with more than 800 amino acids collected from both the CASP15 and CAMEO datasets. The average protein length in this set is 1239 amino acids.

| ProDESIGN-LE | ABACUS-R | Protein<br>MPNN_002 | Protein<br>MPNN_020 | ESM-IF | CarbonDesign |
| --- | --- | --- | --- | --- | --- |
| 36.4% | 36.3% | 46.9% | 41.8% | 32.6% | **55.1%** |

**Table S6:** Evaluation on the testing set of orphan proteins measured with sequence recovery rate.

| ProDESIGN-LE | ABACUS-R | Protein<br>MPNN.002 | Protein<br>MPNN.020 | ESM-IF | CarbonDesign |
| --- | --- | --- | --- | --- | --- |
| 38.9% | 32.6% | 38.5% | 44.3% | 46.2% | **49.1%** |

**Table S7:** Evaluation of CarbonDesign in predicting pathogenicity of variants with the testing set of clinically curated variants in ClinVar.

| Methods | BRCA1 | PTEN | TP53 | MSH2 | average |
| --- | --- | --- | --- | --- | --- |
| ESM-1v | 0.896 | **1.000** | **0.994** | 0.812 | 0.926 |
| ProGen2 | 0.876 | **1.000** | 0.952 | **0.844** | 0.918 |
| CarbonDesign | **0.933** | 0.986 | 0.984 | 0.822 | **0.931** |

**Table S8:** Evaluation on *de novo* backbone structures from RFDiffusion at varying noise levels, measured using scTM score.

| Methods | Length<br>200 | Length<br>300 | Length<br>400 | Length<br>500 | Length<br>600 |
| --- | --- | --- | --- | --- | --- |
| **CarbonDesign(small noise)** | 0.84 | 0.69 | 0.58 | 0.58 | 0.48 |
| **CarbonDesign(high noise)** | **0.89** | **0.80** | **0.74** | **0.64** | **0.54** |

**Table S9:**
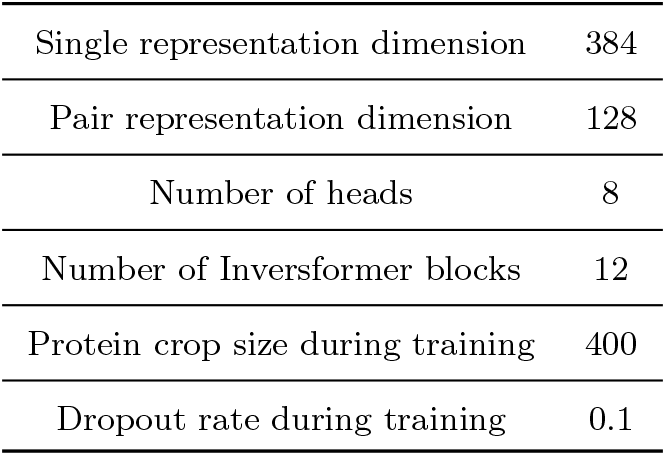
Hyperparameters of CarbonDesgin architecture.

